# Emergence of High-Order Functional Hubs in the Human Brain

**DOI:** 10.1101/2023.02.10.528083

**Authors:** Fernando A.N. Santos, Prejaas K.B. Tewarie, Pierre Baudot, Antonio Luchicchi, Danillo Barros de Souza, Guillaume Girier, Ana P. Milan, Tommy Broeders, Eduarda G.Z. Centeno, Rodrigo Cofre, Fernando E Rosas, Davide Carone, James Kennedy, Cornelis J. Stam, Arjan Hillebrand, Mathieu Desroches, Serafim Rodrigues, Menno Schoonheim, Linda Douw, Rick Quax

**Affiliations:** Dutch Institute for Emergent Phenomena (DIEP), 1090 GL Amsterdam, The Netherlands; Institute for Advanced Study, University of Amsterdam, Amsterdam, 1012 GC, the Netherlands; Korteweg de Vries Institute for Mathematics, University of Amsterdam, Science Park 105-107, 1098 XG Amsterdam, the Netherlands; Amsterdam UMC location Vrije Universiteit Amsterdam, Anatomy and Neurosciences, Boelelaan 1117, Amsterdam, The Netherlands; Clinical Neurophysiology Group, University of Twente, Twente, The Netherlands; Sir Peter Mansfield Imaging Center, School of Physics, University of Nottingham, United Kingdom; Median Technologies, 1800 Route des Crêtes, 06560 Valbonne, France; MCEN Team, Basque Center for Applied Mathematics, Bilbao, Bizkaia, Spain; MathNeuro Project-Team, Inria at Université Côte d’Azur, Sophia Antipolis, France; Department of Clinical Neurophysiology and MEG Center, Amsterdam Neuroscience, Vrije Universiteit Amsterdam, Amsterdam UMC, Amsterdam, The Netherlands; Departamento de Electromagnetismo y Física de la Materia and Instituto Carlos I de Física Teórica y Computacional, Universidad de Granada, 18071, Granada, Spain; Institut Des Maladies Neurodégénératives, UMR 5293, Université de Bordeaux, CNRS, Bordeaux Neurocampus, 146 Rue Léo Saignat, 33000, Bordeaux, France; CIMFAV-Ingemat, Facultad de Ingeniería, Universidad de Valparaíso, Valparaíso, Chile; Institute of Neuroscience (NeuroPSI), Paris-Saclay University, Centre National de la Recherche Scientifique (CNRS), Gif-sur-Yvette, France; Centre for Complexity Science, Imperial College London, United Kingdom; Radcliffe Department of Medicine (RDM), University of Oxford - UK; Ikerbasque, The Basque Foundation for Science, Bilbao, Bizkaia, Spain; Computational Science Lab, University of Amsterdam, Amsterdam, 1098 XH, The Netherlands

## Abstract

Network theory is often based on pairwise relationships between nodes, which is not necessarily realistic for modeling complex systems. Importantly, it does not accurately capture non-pairwise interactions in the human brain, often considered one of the most complex systems. In this work, we develop a multivariate signal processing pipeline to build high-order networks from time series and apply it to resting-state functional magnetic resonance imaging (fMRI) signals to characterize high-order communication between brain regions. We also propose connectivity and signal processing rules for building uniform hypergraphs and argue that each multivariate interdependence metric could define weights in a hypergraph. As a proof of concept, we investigate the most relevant three-point interactions in the human brain by searching for high-order “hubs” in a cohort of 100 individuals from the Human Connectome Project. We find that, for each choice of multivariate interdependence, the high-order hubs are compatible with distinct systems in the brain. Additionally, the high-order functional brain networks exhibit simultaneous integration and segregation patterns qualitatively observable from their high-order hubs. Our work hereby introduces a promising heuristic route for hypergraph representation of brain activity and opens up exciting avenues for further research in high-order network neuroscience and complex systems.

## 1. INTRODUCTION

### Context

Networks offer a universal framework to encode information about interactions in a complex system, which often involve three or more sub-systems in entangled ways. In standard network approaches, high-order interactions are often approximated via pairwise interactions, whereby 3-point interactions between nodes A, B, and C within a network are inferred through the existence of a clique, i.e., pairwise interactions linking A and B, B and C, and C and A. Such approximations, while reasonable, may not be able to capture all important aspects of the interactions. For instance, communication networks can exhibit scenarios where communication between pairs does not necessarily imply simultaneous communication across the three instances (A, B, C) [1].

The inclusion of high-order interactions and connectivity in the study of networks can be seen as a more realistic and informative way of modeling complex systems. However, this induces an important combinatorial burden. To give an idea, in a network with N nodes, one has up to 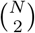 possible pairwise interactions, up to 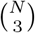 possible three-point interactions, and more generally, up to 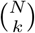, (k ≤ N) possible interactions between k-nodes in a high-order network. While considering the high-order structure of complex systems brings with it a host of combinatorial challenges, it is nonetheless essential for a more realistic modeling and enhanced understanding of complex systems.

In this vein, there has been a significant effort recently in quantifying high-order interactions in multiple complex systems [1–4], from ecology [5] to social contagion [6, 7], across different time and spatial scales [8–10], and for areas such as the synchronization of coupled oscillators [11], gene expression [12] and psychometrics [13] — just to name a few. While some of these approaches focus on the topological analysis of high-order networks [2], others use information theory for inferring high-order statistical interdependence in complex systems, including the brain [8, 14–18].

In this work, we attempt to bridge topological and information-theoretic approaches and build a mathematically well-grounded high-order connectivity framework using high-order interdependencies. More specifically, the framework presented here advances the methodology on how to quantify high-order interactions and build a hypergraph representation of the interdependencies observed in time series data.

### Towards high-order brain networks

We apply the methodology developed in this work to the brain — one of the most complex systems [19] — in order to build high-order representations of functional brain activity from time series recorded from 92 brain regions using resting state functional Magnetic Resonance Imaging (rs-fMRI). Indeed, considering the brain as a network [20–23] offers ubiquitous ways to encode information about the interactions between different areas of the brain at different spatial and temporal scales and for different imaging modalities.

Classical research in neuroscience has led to a microscopic view that neuronal architecture is based on pairwise interactions due to synaptic dyadic connectivity between neurons. Indeed, dendrites of post-synaptic neurons have synapses with numerous pre-synaptic neurons (up to thousands) [24]. As a consequence, mesoscopic and macroscopic brain studies have inherited this dyadic viewpoint. For example, at the macroscopic scale and at the functional level, brain network interactions are typically represented through similarity measures between time series of activity at two brain regions. There is a plethora of ways to define such pairwise interactions, such as the Pearson correlation coefficient, covariance and mutual information. Some of these metrics are tailor-made for neuroscience [25, 26], others for complex systems in general [27]. Despite being reductionist, this dyadic interaction has enabled several breakthroughs in neuroscience.

However, recent neuroscientific observations show that astrocytes, which cover between 105 synapses in mice and 2 × 10^6^ in humans, regulate structural and synaptic plasticity [28]. Crucially, astrocytes regulate hetero-synaptic interactions and the heterogeneity of synaptic strengths, thus enabling higher-order interactions [29]. This induces a shift of paradigm and forces us to rethink our standard theories of neural interactions towards incorporating higher-order structures. We illustrate this high-order paradigm for the brain in Fig. 1. Such a consideration will influence existing brain studies that propose a functional modular architecture and community structure for the brain [30], where pairs of nodes in a brain network act together to execute a specific function. As we propose in this paper, such a modular architecture and community structure also holds beyond dyadic interactions, at the level of high-level interactions.

**FIG. 1:**
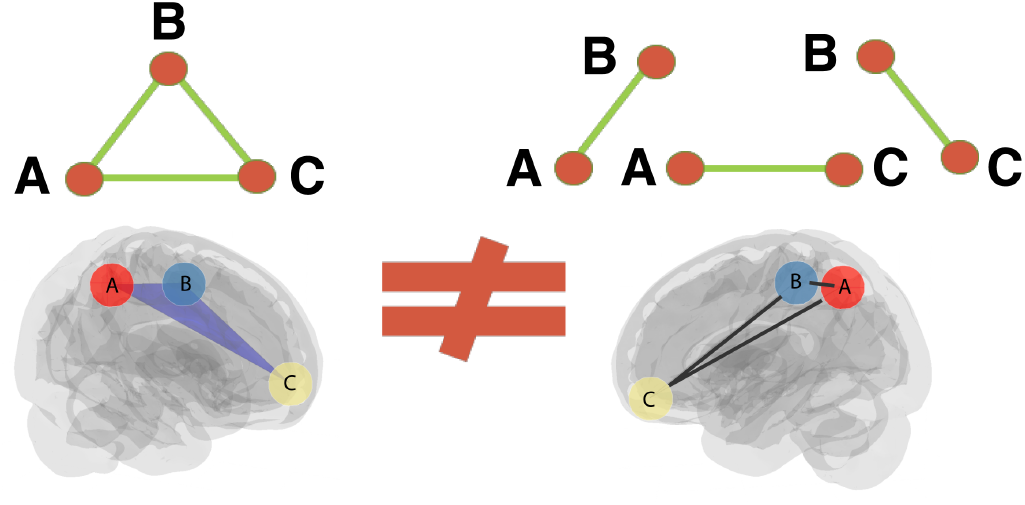
High-order interactions in complex systems, especially in functional brain networks, are often approximated and misrepresented as the sum of pairwise interactions.

### Proposed approach

Building a high-order network from time series requires multivariate signal processing and the systematic identification of its most relevant high-order edges. With a suitable high-order connectivity rule, one can define high-order hubs in a network, which circumvents the combinatorial complexity issue by enumerating only the most important high-order edges without prior knowledge of what the most important hyperedges are.

Fig. 2 sketches our proposed multivariate signal processing methodology to construct high-order networks. Our method relies on heuristics used in the past for pairwise interactions [31] that enable us to define high-order networks as uniform hypergraphs [32]. In pairwise interactions, one defines edges via any reliable pairwise similarity metrics between two nodes in a network. Similarly, this work relies on multivariate similarity metrics to define high-order weights for hyperedges with fixed size, i.e., in a uniform hypergraph. We further utilize the fact that a uniform hypergraph can be represented as a high-order adjacency matrix, accounting for the connectivity between hyperedges [33]. Consequently, multiple algorithms already used for pairwise adjacency matrices such as Eigenvector centrality, modularity, or betweenness centrality could be inherited in the high-order context [33, 34], speeding up the methodological bridge from pairwise to high-order networks.

**FIG. 2:**
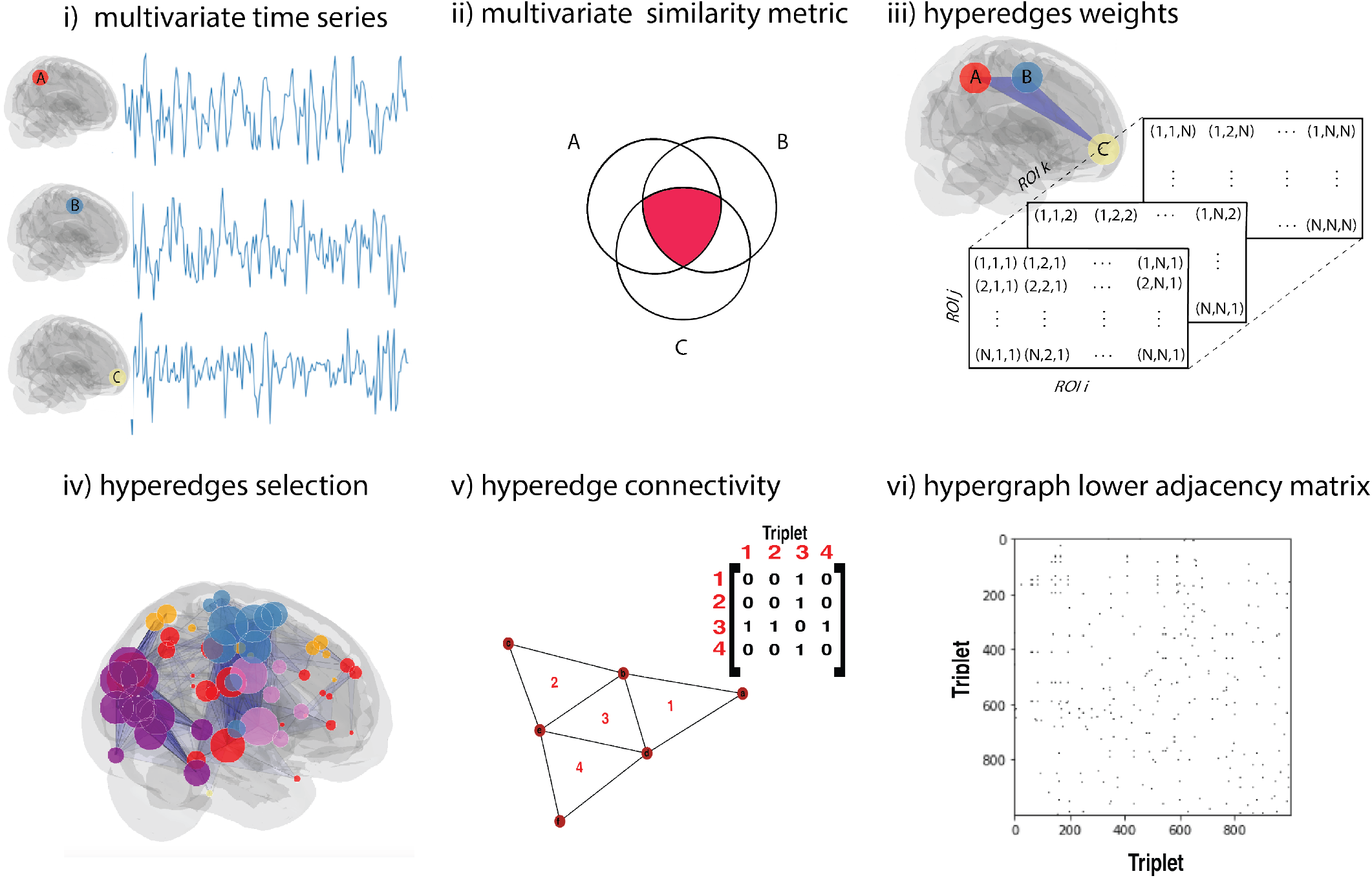
Heuristic construction of a uniform hypergraph: i) We start with multivariate times series as inputs, which in this work are BOLD signals from resting-state fMRI. ii) In analogy with the pairwise case, we define high-order connectivity weights via multivariate statistical dependencies estimations. High-order statistical dependencies are rigorously quantified via, e.g., Multivariate Interaction Information or Total Correlation (see appendix). iii) Once hyperedges’ weights are defined, iv) we can explore different ways to select the most important hyperedges. v) We can then also explore high-order connectivity rules to iv) represent the hypergraph as an adjacency matrix. Consequently, each high-order statistical similarity measure could potentially define a uniform hypergraph from time series [36, 37].

We demonstrate the performance of our proposed methodology on neuroimaging data from the Human Connectome Project (HCP) [35], focusing on the brain data of a cohort of 100 individuals in resting state condition. We use time series corresponding to the resting-state BOLD brain activity of 92 regions of the AAL atlas [38], which yields up to 125, 580 3-point interactions. Thus, despite our new methodology’s advantage, i.e., brain activity can be represented via a high-order adjacency matrix, the combinatorial complexity of such a hypergraph requires that we focus our study on the most important hyperedges. Following the notions from network science, these hyperedges are the most relevant nodes in a network, i.e. the hubs. This leads us to investigate high-order hubs in brain networks via the Eigenvector centrality of the hypergraph.

Our analyses reveal that those high-order hubs are far from random: As we show, each multivariate similarity metric is compatible with a different system in the brain. For instance, we find that some three-point hubs provide segregation and integration triplets that are centered within the somatosensory and visual systems. This finding suggests that high-order hubs can be considered emergent high-order properties of functional brain networks. Furthermore, our illustrative example shows that the functional connectivity of a high-order hub centered in the somatosensory system correlates with the gait speed of the individuals.

These findings provide evidence of the broad applicability and relevance of our methodology, suggesting a promising route for hypergraph representation of brain activity — opening exciting avenues for further research in high-order network neuroscience and complex systems.

## 2. METHODOLOGY

This section introduces our methodology to represent the statistical structure of time series as hypergraphs, following heuristics analogous to early developments of pairwise connectivity [39, 40]. Our methodology for general multivariate time series is implemented in five steps, as described in the following (see Fig. 2):

1. We use a given set of N times series as input. As a first step, we assign one node per time series, which will be the basis for building a hypergraph. For time series corresponding to resting-state fMRI, each node corresponds to a different brain region.
2. We choo se an order k *∈ {*3, …, N *}*, and calculate the 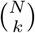 k-order terms associated with a high-order interdependency metric between all possible groups of k different nodes. As in the case of pairwise connectivity, different multivariate similarity measures can be considered, and the optimal choice is usually domain- and problem-dependent. Among the various multivariate information metrics available in the literature [36], in this paper, we focus on two information-theoretic measures to showcase our methodology: Interaction Information (II), which quantifies single tuple statistical dependencies, and Total Correlation (TC), which quantifies cumulated total statistical dependencies over all tuples subsets [1](see Appendix for details).
3. Once all the interdependencies have been calculated, we proceed to hyperedge selection. This is done because the number of k-order interactions among N nodes grows with N ^*k*^, and we would otherwise have a complete weighted hypergraph. There are multiple ways to access the significance of edges in a network, and those strategies also translate to high-order connectivity. Here, for simplicity, we threshold the high-order interdependencies and keep the strongest hyperedges in the hypergraph.
4. We follow a procedure to encode the resulting k-uniform hypergraph as an “hyper-adjacency matrix.” Among the multiple alternatives [33], the one used in this work inherits the lower adjacency matrix representation of simplicial complexes into uniform hypergraphs, which was also recently adapted to develop vector centralities on hypergraphs [41]. Put simply, two hyperedges of dimension k are connected if they share an hyperedge of dimension k *–* 1 [33, 34, 42]. For example, two triangles are connected if they have an edge in common.
5. As a final step, after our multivariate signal processing and the hyper-adjacency matrix has been specified, many topological features from network science can be leveraged. In this work, we use an extension of the Eigenvector centrality, introduced in [42], to investigate high-order hubs in the human brain. High-order hubs were explored recently in network science in diverse settings [33, 43]. In this work, we will rely on the Perron-Frobenius theorem [44] to explore high-order hubs of uniform hypergraphs. Hence, we investigate the spectra of the hyper-adjancency matrix and identify hubs as triplets with higher eigenvector centrality. Representing the hypergraph as an hyper-adjacency matrix lets us, in turn, also calculate other features, such as modularity and betweenness centralitiy, which can be explored in different contexts.

Please note that the combinatorial complexity of k-order hyperedges over a network of N nodes is *𝒪* (N ^*k*^). Our approach is to circumvent this important computational bottleneck by relying on network analyses on the hyper-adjacency matrix. We illustrate this procedure for a simple network of triangles in Fig. 2 v). This methodology allows us to infer realistic *k* -body interactions from time series and represent them as a hypergraph, which is crucial for the theoretical development of network science.

## 3. APPLICATION TO RS-FMRI TIME SERIES

As a proof of concept, we apply the methodology described above to fMRI data from 100 unrelated individuals (young adults) from the Human Connectome Project [35] and report group averaged higher-order inter-dependency results. To illustrate the flexibility of our framework, we showcase our methodology using two multivariate information metrics, Interaction Information (II) and Total Correlation (TC) (see appendix for details), that correspond to alternative generalizations of multivariate correlation coefficients. For both metrics, we selected the 1000 strongest triplets in the hypergraph, which allowed us to construct an adjacency matrix representation of the hypergraph.

### Interaction Information and the somatosensory system in the brain

The hyperedges denfined via the Interaction Information (II) are illustrated in Fig. 2 via the projection of the corresponding triplets (iv) and the adjacency matrix (vi) using data visualization methods for simplicial complexes developed in [45, 46].

We find that the strongest triplets are consistent with the network organization of the brain in sub-networks, and we follow [47] for the color code.

Fig. 3(a) shows the distribution of the Eigenvector Centrality (EC) of the 1000 triplets with the highest II. From this histogram, it is clear that only a small fraction have an EC far from zero (only 42 triplets are highlighted in the red circle). Given the combinatorial complexity of high-order interactions, the interpretation of the whole of high-order interactions in brain networks will become more evident if one projects only the triplets with higher high-order EC in the brain (Fig 3 a). Overall, for II, we observe that all central triplets with the highest EC share two nodes and one link in common, covering the whole brain, yielding a pattern that we identify here as “high-order hubs.”

**FIG. 3:**
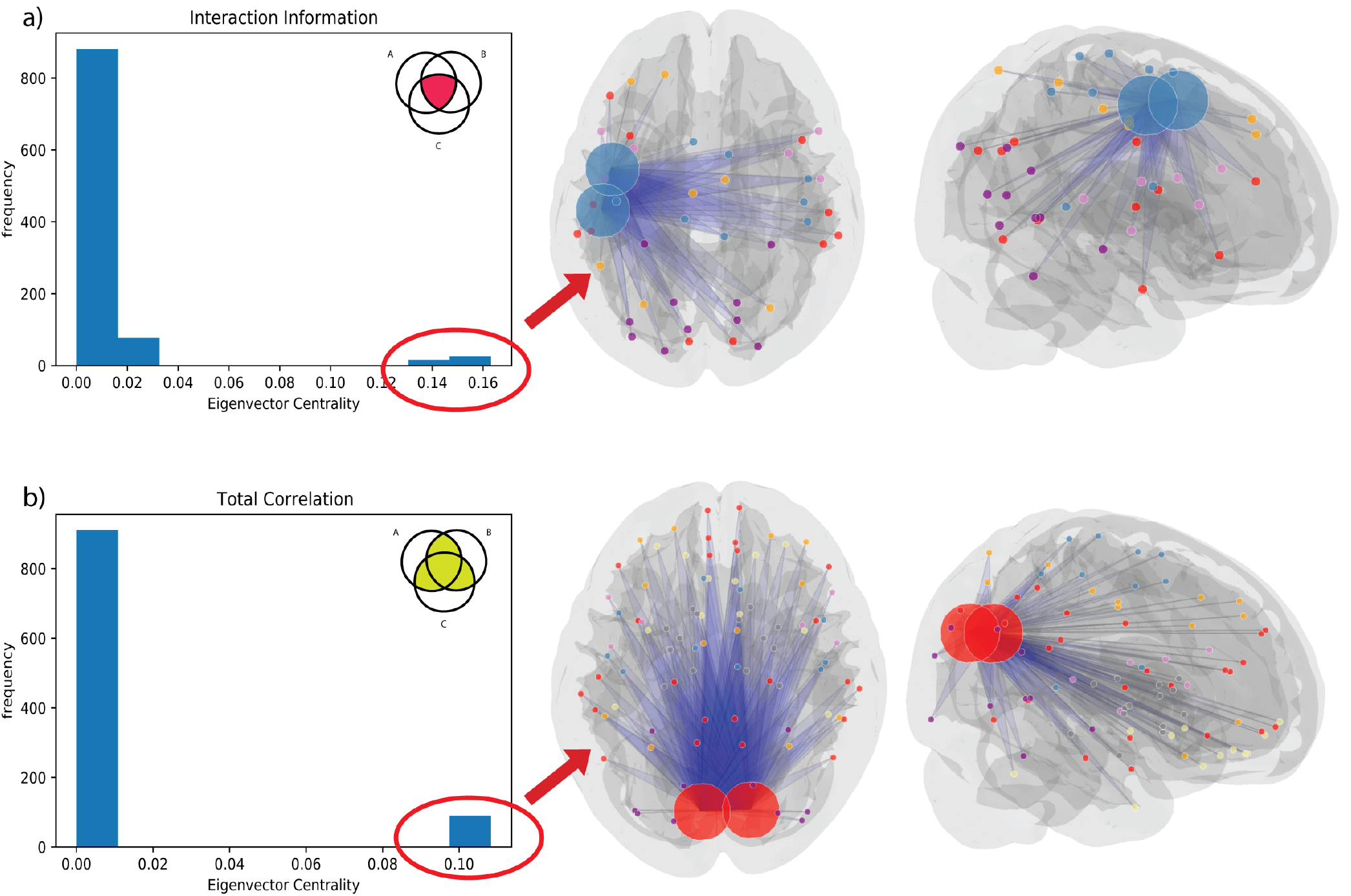
Emergence of high-order hubs in functional brain networks: When computing the high-order EC based on the 1000 strongest triplets both for Interaction Info (a) and total correlation (b), only a small fraction of those triplets have non-zero centrality. Projecting triplets with higher EC unveils the emergence of a high-order hub centered in the sensory-motor system for II (a) and visual system (b).

The two nodes that are shared between the 42 triplets with high EC shareare the pre- and post-central gyri. As we discuss below, given the role of those two areas in the functioning of the brain [48, 49], we associate those emergent high-order hubs with the primary sensory-motor system. One interesting feature about the pre- and post-central gyri is that they are “topographically organized”, i.e., adjacent areas of the motor (or receptive) surface of the human body are mapped in neighboring areas of the pre- (and post-)central gyrus. The pre- (and post-)central gyrus corresponds to the primary motor (and somatosensory) cortex, and the relative size of the cortical representation reflect the density of motor (and somatosensory) nerves in each body part.

The somatosensory strip (post-central gyrus) contains an inverted somatotopic map of the opposite side of the body that nearly mirrors that of the motor strip (precentral gyrus). Yet, very recently, this map was revisited, and integrative regions of the motor strip were also reported, relating inter-effector regions with prefrontal, insular, and subcortical regions of the Cingulo-opercular network (CON), critical for executive action and physiological control, arousal, and processing of errors and pain [50]. Given that the HCP database consists of mostly right-handed individuals, the fact that the high-order hubs are in the brain’s left hemisphere is consistent with the well-known asymmetry in the functional connectivity of sensory-motor areas due to handedness [51, 52]. Moreover, due to the necessity of sensory-motor integration to execute one movement, activity in those two areas is highly coupled. For instance, when a person moves the right hand, increased blood flow is recorded in the left pre-central gyrus, and simultaneous activation is observed in the corresponding area of the post-central gyrus [53, 54].

Integrating our finding with the current understanding of the pre- and post-central gyrus, one plausible interpretation is that, given that pre- and post-central gyrus are highly coupled in the brain, any relevant interaction between these two areas and another third area in the brain would be at least a three-point interaction between sensory-motor and other cortical regions.

### Total Correlation and the visual system in the brain

Next, we turn to our second multivariate information metric, Total Correlation (TC), to build a 3-hypergraph from rs-fMRI signals. We used a similar procedure to II, i.e., we computed the TC of all possible triplets in the AAL atlas and chose the 1000 triplets with the highest TC to build a uniform hypergraph.

Again, only a small fraction of the triplets have an EC not close to zero. When we project those triplets with high EC in the brain, we find that all triplets involve two specific areas in the brain. In this case, however, the high-order hubs are associated with the visual Broadman area 17, namely the left and right cuneus, as illustrated in Fig. 43 (b). Each cuneus (from the left and right hemispheres) receives visual information via pulvinar thalamus from the contralateral superior retina representing the inferior visual field, as well as via geniculate thalamus. The cuneus is also known for its extraretinal role, and it is modulated by, e.g., attention, working memory, and reward expectation [55–57]. Given the contralateral division of the visual maps of the cuneus and its role in the brain, and similar to the sensory-motor case, under a high-order perspective, the integration or modulation of visual information is likely promoting at least three-point interactions defined by TC.

It is important to notice that our results align with the recent work by Faes *et al*. [18], where the authors reported that the redundant cores are more prevalent in the visual and motor systems. Our work, which builds a hypergraph from inter-dependencies, adds the fact that the most central hyperedges in the hypergraph representation of the brain are compatible with the current knowledge in functional neuroanatomy. At least from a qualitative perspective, the emergent pattern of the redundant high-order hubs defined via II and TC are compatible with local segregation and global integration principles.

### Synergistic high-order hubs in the brain

We illustrated hypergraphs for the strongest, positive II, and TC in the previous section. However, every multivariate information metric or any other measure of statistical interdependence between variables could be used to define hypergraphs from time series. Our choice for II and TC falls in the category of so-called redundant interactions, where high-order interactions exist jointly but can also be split into parts. Yet, another type of interaction, the synergistic interaction, exists only in high-order but not in smaller pieces. Both types of interactions are relevant since redundancy is related to robustness, whereas synergistic interactions are related to integration.

Since synergistic and redundant interactions are complementary, we also show hypergraphs defined and binarized by the strongest negative II. In Fig. 4 (a), we illustrate the hypergraph defined from synergistic II following a similar procedure as in previous sections, but the only difference is that we selected the 1000 most negative triplets for the hypergraph. An interesting feature of those hubs is the emergence of cortical-sub-cortical interactions mediated by areas from the Default Mode Network, especially the angular gyrus, as illustrated in Fig 4(a).

**FIG. 4:**
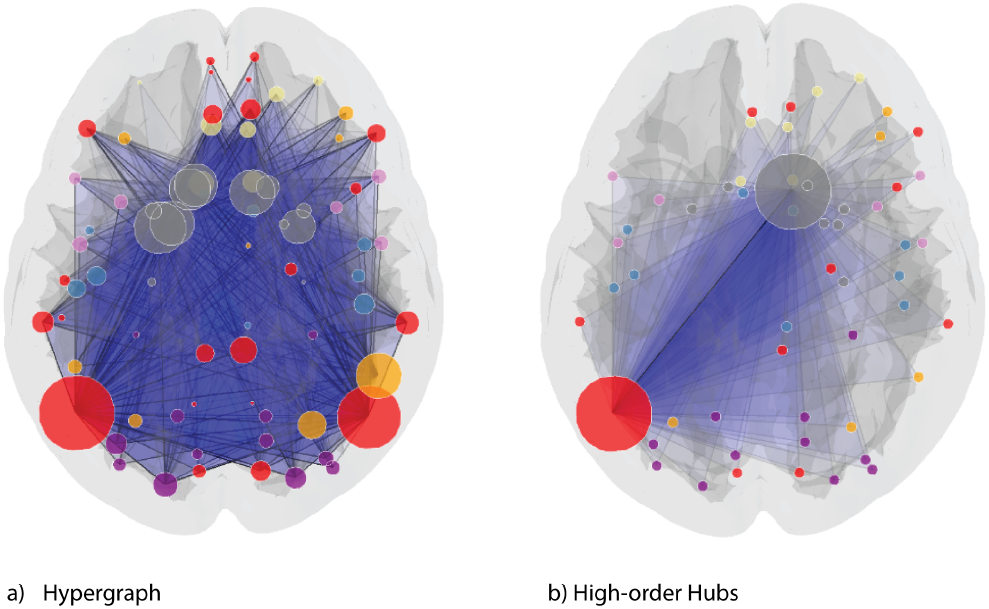
(a) Hypergraph build from 1,000 triplets with high synergy illustrates the integration between default mode network (red nodes) and sub-cortical regions (gray nodes). (b) In this case, synergistic high-order hubs are centered on the left angular gyrus and the right nucleus accumbens.

The angular gyrus is considered a higher-order associative cortical region relevant for integrating multiple sensory systems, language comprehension, number processing, spatial attention, and memory retrieval [58]. Recently, a cross-species similarity study comparing humans and macaques shows a gradient of evolutionary change where most pronounced changes are related to areas of the Default Mode Network, especially the angular gyrus [59]. Therefore, the fact that the synergistic hypergraph is related to the angular gyrus is also compatible with recent findings relating synergy to evolution and cognition [18]. More specificaly, the high order-hubs that are the left angular gyrus and right nucleus accumbens, as illustrated in Fig. 4(b).

### Paradigmatic evidence of the relevance of high-order hubs in neuroscience

In the previous sections, we have introduced a heuristic methodology to build uniform hypergraphs from brain signals and the most important high-order hubs in this framework. One current challenge that exists in clinical network neuroscience is whether one can find a relationship between large-scale brain networks and behavioural outcomes [60]. The relationship between functional connectivity and behavior is complex and not well understood, and as a result, the R-squared value of this relationship is often low. Yet, other factors beyond brain connectivity also contribute to individual differences in behavior.

Thus, one question that naturally emerges in this context is whether the uniform hypergraphs introduced in this work would contribute to understanding this quest. Given that the high-order hubs observed here using redundant II were associated with the pre and post-central gyrus, we naively raised the question of whether any motor behavioural trait could correlate with the strength of these high-order hubs.

To this aim, we investigate whether the II of one of the 42 high-order hubs associated with motor areas would correlate with the annotated gait speed of the 100 unrelated healthy individuals analysed in this work. Gait speed is the speed at which participants walk a short distance at their usual pace. It can be used as a clinical indicator of mobility [61] and overall health in different populations [62]. We found that the strength of one specific triplet with high eigenvector centrality, constituted of the left pre and post-central gyrus and the right temporal lobe, has significant correlation (*p ≈* 0.0007, *R*^2^ = 0.113) with the gait speed of the individuals, as illustrated in Fig. 5.

**FIG. 5:**
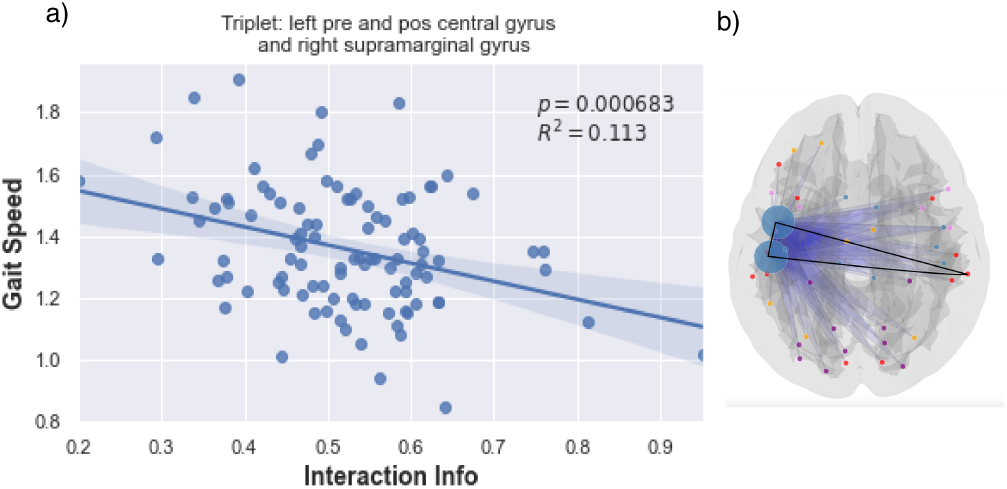
(a) Interaction Information (II) of the triplet defined by the left pre- and post-central gyrus and right supramarginal gyrus correlates significantly with the gait speed of the individuals. This triplet is highlighted in black in the righ panel (b).

## 4. DISCUSSION

Despite the numerous ways of creating networks across domains, there is no established methodology to build hypergraphs from empirical data in network science. In this work, we propose a methodology to build hyper-graphs based on high-order inter-dependencies of time series data, followed by simple connectivity rules for defining hyperedges in a high-order network – namely a uniform hypergraph. In contrast to general hypergraphs that need an incidence matrix to be represented, uniform hypergraphs can be represented as a simple lower adjacency matrix that inherits the most common algorithms in network science. By leveraging this opportunity, our approach opens the possibility to a plethora of new applications using established network algorithms applied to high-order interdependencies. This methodology can be seen as a natural extension of pairwise connectivity for high-order interactions based on high-order statistical interdependencies.

To showcase our methodology’s capabilities, we applied it to analyse brain networks – more specifically, rs-fMRI time series of 100 individuals from the Human Connectome Project. When projecting the resulting high-order hubs into a brain atlas, we observed the emergence of high-order hubs compatible with well-known systems in the brain. We applied our method using two different similarity measures, Interaction Information (II) and Total Correlation (TC), resulted in high-order hubs compatible with the sensory-motor and visual systems, respectively.

Our results are a promising starting point for advancing the hypergraph representation of complex data, but they are far from complete. Further developments of high-order networks face similar limitations and already existing choices for functional connectivity [21]. In addition, there is an inherent issue related to the combinatorial complexity that grows with the number of high-order interactions included. We addressed this combinatorial problem by studying the Eigenvector Centrality of the hyper-links. Due to the fact that uniform hypergraphs have adjacency representation, one can use standard network algorithms for this aim. This approach significantly decreased the complexity and number of hyperedges in our work.

Future developments based on our work are at least two-fold. From a methodological point of view, given that an intermediate output of our work is a high-order adjacency matrix, in principle, any metric built upon an adjacency matrix in the pairwise case has a high-order counterpart. Thus, one can study multiple high-order metrics beyond high-order hubs, such as clustering, betweenness centrality, random walks, and modularity, to name a few. Furthermore, in our work, we illustrated possible ways of building high-order networks from empirical data that could be further explored in different contexts. We focused here on some multivariate information theory measures, but in principle, any multivariate similarity metric can be used to build uniform hyper-graphs (see, e.g., [9]). The chosen measures here quantify statistical dependencies and generalise correlations, and are both central probabilistic and topological measures, and hence provide straightforward tools to define and quantify neural assemblies and extend the correlation theory of the brain to higher order statistical interactions (see, e.g., [63] for an introduction to neural assemblies, and [64] for previous topological investigations).

From a brain imaging perspective, tailoring this approach to other image modalities, such as EEG, MEG, and DTI, is also of great interest. From a clinical neuroscience perspective, much work has been done trying to correlate pairwise interactions and behavioral traits at individual and group levels, and correlation is at the basis both of synaptic learning rules, and perceptual binding of multiple features into a single coherent emergent perceived object [65, 66]. Since our work gives strong evidence that some systems in the brain may act as a high-order network, this work opens a perspective of correlating high-order properties of brain networks with specific clinical traits, as was evidenced here via gait speed. Various aspects of cognition and behavior are hypothesized to be emergent and high-order phenomena [67]. Therefore, further application in large-scale databases are necessary to unveil the relations between high-order hubs, other high-order centrality metrics, brain imaging andhuman behaviour. Ultimately, since this work is built on multivariate time series signals, these ideas could be translated more generally to other complex systems.

## Supporting information

Interaction Information - High-order sensorimotor hub

## ACKNOWLEDGEMENTS

This work is supported by the Dutch Institute for Emergent Phenomena (DIEP) cluster at the University of Amsterdam. This work was developed during the research fellowship “High-Order Interactions in Complex Systems” awarded to F. A. N. Santos by the Institute for Advanced Studies at the Universit of Amsterdam. F. A. Santos would like to acknowledge research visits at the The Abdus Salam International Centre for Theoretical Physics (ICTP) in Trieste. F. A. N. Santos would like to thank fruitful discussions with Clelia de Mulatier, Giovanni Petri, Rikkert Hindriks, Peter Sloot, Piet Boumann, Matteo Marsilli, Jeroen Geurts, Tiziano Squartini, and J. P. Crutchfield during different stages of this work. SR acknowledges support from Ikerbasque (The Basque Foundation for Science), the Basque Government through the BERC 2022-2025 program and by the Ministry of Science and Innovation: BCAM Severo Ochoa accreditation CEX2021-001142-S / MICIN / AEI / 10.13039/501100011033 and through project RTI2018-093860-B-C21 funded by (AEI/FEDER, UE) and acronym “MathNEURO”. MD and SR acknowledge the support of Inria via the Associated Team “NeuroTransSF”. RC acknowledge the support the Human Brain Project, H2020-945539.

## APPENDIX

### Multivariate information and high-order statistical dependencies measures

Here, we provide the mathematical details on the computation of Multivariate Information theory measures crucial for defining weights in our high-order brain networks. On a pairwise level, Information quantities are well understood and defined by Shannon [68] and Kullback [69]. The Shannon entropy of a single random variable *X, H*(*X*), with probability distribution *p*(*x*), is defined as follows:

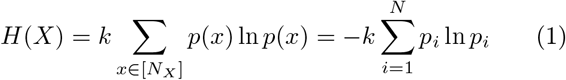

where [N_*X*_] = {1, …, N_*X*_} denotes the codomain or alphabet of X and k = *−*1/ ln 2. It quantifies the amount of uncertainty that is present in the probability distribution. The Shannon entropy is maximum for the uniform probability distribution, and it is zero and minimal for the delta probability distribution concentrated in one value.

The joint entropy is defined for any joint-product of n random variables 𝕏 = (*X*_1_, …, *X*_*n*_) and for a probability joint-distribution p(x_1_, …, x_*n*_) by

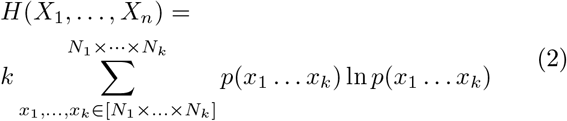

where [*N*_1_ *× … × N*_*k*_] = {1, …, *N*_*j*_ *× … × N*_*k*_} denotes the codomain (also called alphabet) of (X_1_, …, X_*k*_).

The amount of shared information between 2 variables, is given by the mutual information as follows [70]:

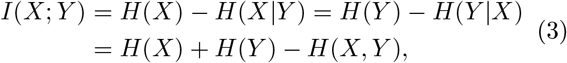

where *H*(*X* |*Y*) is the conditional entropy, which is given by:

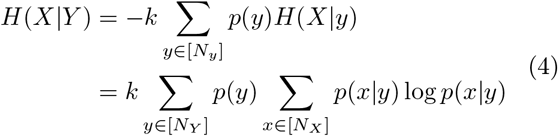

Mutual information measures statistical dependencies in the sense that it is equivalent to say that *X* and *Y* are statistically independent or to say *I*(*X*; *Y*) = 0. It generalises Pearson correlation coefficient *C*(*X*; *Y*) and covariance, and notably we have *I*(*X*; *Y*) = 0 implies *C*(*X*; *Y*) = 0 but not the converse.

When we move on to measuring high-order relations, a variety of measures have been proposed [1, 4, 14, 71, 72]. In this work, the method to construct high-order networks based on the definition of uniform hypergraphs could be used in principle with any of those high-order measures as a candidate for high-order similarity measures. In this work, we focus on the two main multivariate information measures that quantify statistical dependencies, namely Interaction Information (II) and Total Correlation (TC). Indeed, there is essentially two different generalisations of pairwise Mutual information to the higher order multivariate case: Multivariate Mutual-information or Interaction Information (II), [73] and Total Correlation (TC), [71] such that the single variable case is entropy and the bivariate case is pairwise MI. The multivariate MI can be written as an expansion of the entropies on the variables:

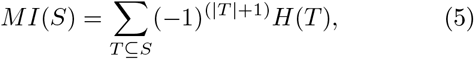

where *S* = {*X*_1_, *X*_2_, … *X*_*N*_ }, and the sum runs over all *T* subsets of *S* and |*T*|denotes the cardinal of *T*. For example, we have:

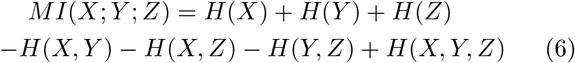

Similarly the multivariate II for *N* number of variables has the following expansion (here we use the sign convention of [74]):

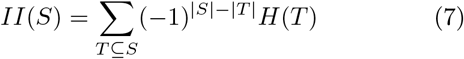

For three variables the interaction information *II*(*X, Y, Z*) can be written as follows:

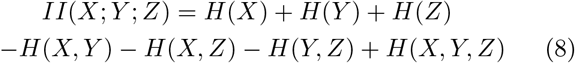

Notably, entropies are just sums of II:

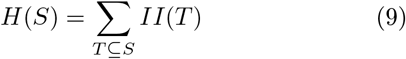

The second metric explored in this work was TC [71], which quantifies the redundancy or dependency among the variables in the set, which is defined as :

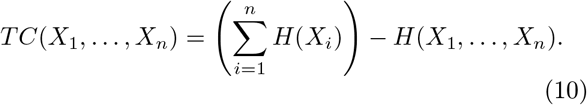

It is equal to the Kullback–Leibler divergence between the joint distribution and its marginals, and hence always positive or nul.

While II and MI are essentially the same, as they only differ by an alternate sign convention :

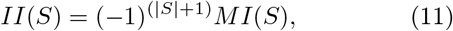

TC is quite different and quantifies the total amount of II, or the “generalised correlation”, in the sense that it sums over all possible higher order Interactions Informations (over all pairs, triplets…):

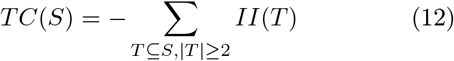

Just as for two variables, TC and II or MI quantify the statistical dependencies among N variables but in a different way :

- For TC: it is equivalent to say that the N variables {*X*_1_, *X*_2_, … *X*_*N*_} = *S* are statistically independent or that *T* (*S*) = 0 [75].
- For II or MI: it is equivalent to say that the N variables {*X*_1_, *X*_2_, … *X*_*N*_} = *S* are statistically independent or that *II*(*T*) = 0 for all subsets T of S, with |*T* | *≥* 2 [1].

In other terms, while TC quantifies global statistical dependencies, II provides a refined local definition of statistical dependencies.

#### Synergy and Redundancy

An important specificity of II and MI with respect to TC is that they can be negative for (*N ≥* 3), a phenomenon called generically synergy [76]. Synergistic sets of random variables only emerge in high-order (*N ≥* 3), but not in pairwise interactions. In this work, we considered as synergistic information the negative Multivariate Interaction Information, while the positive Multivariate II are considered redundant following topological and pseudo-convexity (sub-harmonicity) arguments of [1]. Remarkably those synergistic interactions between N variables cannot be approximated via pairwise interactions or by other k interactions with *k* < *n* [1, 77]. This motivates to consider them as intrinsically emergent collective interactions.

Notice that all Multivariate Info metrics discussed in this appendix can be chosen as k-body similarity metric, thus allowing the construction of uniform hypergraphs from interdependencies. However, in other to build more complex structures (distinct form uniform hypergraphs) further work is required, potentially based on partial information decomposition or information topology, that would allow us to build more general hypergraphs, as well as simplicial complexes.

#### Uniform hypergraph representation of High-order Inter-dependencies

In this work, our goal is to build binary uniform hypergraphs from the strongest redundant or synergistic high-order (multivariate) inter-dependencies. This approach bridges high-order interactions and empirical complex systems more simpler since uniform hypergraphs have adjacency matrix representation [32]. To do so, we have to use the following definitions. Roughly speaking, a hyper-graph is a generalization of a graph where edges can join any number of vertices. For standard graphs, however, edges are connected only between two vertices. An undirected hypergraph *H* is defined as the pair *H* = (*V, HE*) where *V* is the set of elements called nodes or vertices and *HE* is a set of subsets of *V* called hyperedges. Multiple complex systems could benefit from being represented as a hypergraph or other high-order representations [78].

Especially in the brain, where the order of 100 billion neurons and 100 trillion synapses [79], among other factors contributes to the expectations that its interactions are better represented if it is encoded as a network in a high-order manner. On the other hand, hypergraph (or any other high-order representation of complex systems) also brings a combinatorial complexity that needs to be addressed in this work. For practical purposes, even for the simplest case of a hypergraph made of triplets, in the AAL atlas, we have 125,580 triplets, which makes analysis and interpretation unfeasible without further analysis, which allow us to narrow to the most important triplets.

#### Hyperedge adjacency representation

We say that two hyperedges of size k are adjacent, i.e., connected, if they share a hyperedge of size *k −* 1. Notice that for a given hypergraph or simplicial complex, one could have different ways to define such connectivity, e.g., by sharing edges of smaller sizes and so on [33]. This representation was constructed initially for simplicial complexes [33]

In this work, as we only explored 3-point interactions in rs-fMRI data, the lower adjacency matrix coincided with, and further explored recently for hypergraphs [43]. Formally, the hyperedge adjacency representation can be defined as follows [33, 34]: We say that two *k*-edges *e, f* are *lower adjacent* if there is a *k −* 1-edge *l* such that *l ⊂ e, f* and we denote 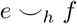, or simply 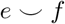.

The *lower adjacency matrix* is defined by

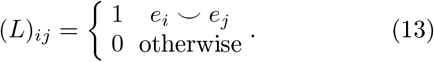

Also, we say that they are upper adjacent if there is a (k + 1)-edge *h* such that *h ⊂ e, f* and we denote by 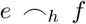, or simply 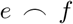. Also, the *Upper adjacency matrix* is defined by

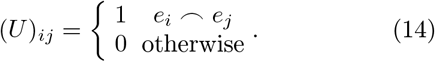

The *Adjacency matrix* for k-edges is defined as

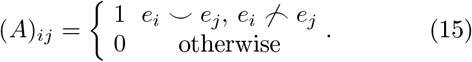

In this work, as we focused only on 3-point interactions in rs-fMRI data, the lower adjacency matrix coincided with the adjacency matrix for 3-edges (triplets).

#### High-order Eigenvector centrality

Multiple hypergraph metrics are natural extensions of network metrics [78], as is the case of high-order hubs used in this work. One of the advantages of the Adjacency representation of hypergraphs, in contrast to incidence matrix representation, is that multiple algorithms are inherited from standard network analysis. This is the case for hypergraph eigenvector centrality. In contrast to standard network representation, where one has up to 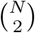 interactions between variables (nodes), in a hypergraph, one has up to 2^*N*^ possible interactions. For each hyperedge size *k*, one has up to 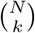 possible interactions. In order to compute High-order eigenvector centrality, one can compute the eigenvector for each hyperedge size k, in the same way, that is computed for networks, as follows.

Let 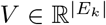, be the eigenvector associated with the highest absolute eigenvalue of one of the adjacency matrix defined in equations (13), (14) or (15), where *E − k* is the cardinally of k-edges. The *eigenvector centrality* of each k-edge in *E*_*k*_ is defined as [32]

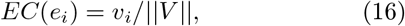

where *v*_*i*_ is the *i − th* entry of *V* and ||.|| is the euclidean norm. Due to the Perron–Frobenius theorem, the eigenvector centrality is unique (up to a constant) and positive.

